# Single-PanIN-seq Unveils that ARID1A Deficiency Promotes Pancreatic Tumorigenesis by Attenuating KRAS Induced Senescence

**DOI:** 10.1101/2020.10.30.361972

**Authors:** Shou Liu, Wenjian Cao, Yichi Niu, Jiayi Luo, Yanhua Zhao, Zhiying Hu, Chenghang Zong

## Abstract

ARID1A is one of the most frequently mutated epigenetic regulators in a wide spectrum of cancers. Recent studies have shown that ARID1A deficiency induces global changes in the epigenetic landscape of enhancers and promoters. These broad and complex effects make it challenging to identify the driving mechanisms of ARID1A deficiency in promoting cancer progression. Here we identified the anti-senescence effect of ARID1A deficiency in the progression of pancreatic intraepithelial neoplasia (PanIN) by profiling the transcriptome of individual PanINs in the mouse model. Interestingly, we found that ARID1A deficiency upregulates the expression of Aldehyde dehydrogenase ALDH1A1, which plays an essential role in attenuating the senescence induced by oncogenic KRAS. Despite that ALDH proteins have been commonly used as cancer stem cell markers, their effect in promoting attenuation of senescence is not known before. Therefore, ALDH proteins could be considered a potential adjuvant drug target in treating pancreatic cancer.

## INTRODUCTION

Pancreatic ductal adenocarcinoma (PDAC) was the third-leading cause of death of cancer in the United States in 2018. It was projected to become the second cause of cancer-related death by 2030 (1). The clear driver mutations of PDAC are *CDKN2A*, *TP53, SMAD4*, and *KRAS*. On the other hand, there are a large number of genes with low-frequency mutations (<5%) (2–4). While these mutations occur at low frequency, they are still statistically significant, which suggests that they play functional roles in promoting tumor development and fitness. From the evolutionary perspective of tumor development, the probability of acquiring one of the mutations in low-frequency-mutation genes is significantly higher than the acquisition of mutations in the very few high-frequency-mutation genes, especially at the very early stage when the prelesion cells are scarce. Therefore, it is reasonable to believe that a portion of the low-frequency mutations can be acquired at the early stage, and they could play essential roles in tumorigenesis.

Among the genes with low-frequency mutations in PDAC, the top hit is *ARID1A* (a subunit of SWI/SNF chromatin remodeling complex) with a 3.6% mutation rate. In addition to PDAC, *ARID1A* is also frequently mutated in other cancer types, including 45.2% of ovarian cancer, 18.7% of gastric cancer, 18.6% of bladder cancer, 13.7% of hepatocellular cancer, 11.5% of melanoma, 9.4% of colorectal cancer, 8.2% of lung cancer and 2.5% of breast cancer (5). ARID1A involves in regulation of many biological processes of cells, including differentiation, proliferation, and apoptosis (6). Therefore, it is greatly desired to study the potentially ubiquitous mechanisms of ARID1A deficiency that facilitate the tumorigenesis of various types of cancers.

Interestingly, in pancreatic cancer, recent studies have shown that ARID1A is necessary to maintain terminal differentiation of pancreatic acinar cells and the knockout results in the accelerated formation of acinar-to-ductal metaplasia (ADM) and neoplastic transformation (7, 8). Meanwhile, Kimura *et al.* and Wang *et al*. also showed that *Arid1a* knockout could promote the formation of PanIN and intraductal papillary mucinous neoplasm (IPMN), which are the most common precursor lesions of PDAC (9, 10). However, despite the clear molecular signature of trans-differentiation in acinar cells, the underlying mechanisms for the acceleration of ARID1A-deficiency-promoted PanIN progression remain elusive.

To dissect the mechanisms that drive the PanIN progression, here we applied to single-cell RNA-seq method to profile the transcriptome of individual early-stage PanIN lesions from *Arid1a* knockout and wild type mice. Our results showed that, beyond complex trans-differentiation, *Arid1a* knockout could effectively reduce Kras-induced senescence in PanIN lesions. It is important to point out that cellular senescence has been shown as the major rate-limiting step in Kras-driven PanIN progression (11). Therefore, with the effective attenuation of senescence, *Arid1a* knockout can achieve a significant acceleration of PanIN progression. Mechanistically, we found that Aldehyde dehydrogenases play an essential role in attenuating the senescence by scavenging the reactive oxygen species (ROS) induced by mutant KRAS.

## RESULTS

### Individual PanIN lesion RNA-seq unveils the potential player contributing to the attenuation of Kras-induced senescence in*Arid1a* knockout mice

To identify the mediators that contribute to ARID1A-deficiency-promoted PanIN progression, we followed the PanIN progression of conditional *Arid1a* knockout mice with mutant KRAS (*Arid1a*^fl/fl^;*LSL-Kras*^G12D/+^;*Ptf1a*^CreERT/+^, A^fl/fl^KC or AKC) and the mice without *Arid1a* knockout (*Arid1a*^+/+^;*LSL-Kras*^G12D/+^;*Ptf1a*^CreERT/+^, A^+/+^KC or KC) (**Fig. S1A**). Consistent with the findings from other groups(7–10), we observed that *Arid1a* knockout facilitates the progression of lesions from ADM to PanIN3 (**Fig. S1B-C**). At as early as the 2-month time point, the percentages of ADM, PanIN-1, and PanIN-2 are 74%, 26%, and 0% respectively in KC mice versus 53%, 47%, and 0% in AKC mice. At the 6-month time point, the percentages of ADM, −1, PanIN-2 and PanIN-3 were 77%, 23%, 0% and 0% respectively in KC mice versus 16%, 74%, 9%, and 0.5% in AKC mice (**Fig. S1C**).

To profile the transcriptome of individual PanIN lesions, here we combined laser capture microdissection (LCM) with a highly sensitive single-cell RNA-seq method (MATQ-seq) developed in our lab (12) (**Figure 1A**). In total, we dissected and profiled 20 lesions from two KC mice and 24 lesions from two AKC mice. We only dissected PanIN-1 and PanIN-2 lesions because their duct-like structures can be easily recognized on the frozen sections for dissection (**Figure S2**). As a result, we detected 18,562±1,840 genes on average for individual PanIN lesions, suggesting the successful transcriptome profiling of individual lesions by MATQ-seq. We observed a clear separation between the lesions dissected from KC and AKC mice in the multidimensional scaling plot (**Figure 1B**), indicating the differential gene expressions between PanINs from KC and AKC mice can be robustly detected by our single-PanIN-seq approach (**Supplementary Table 1**).

**Figure 1.**
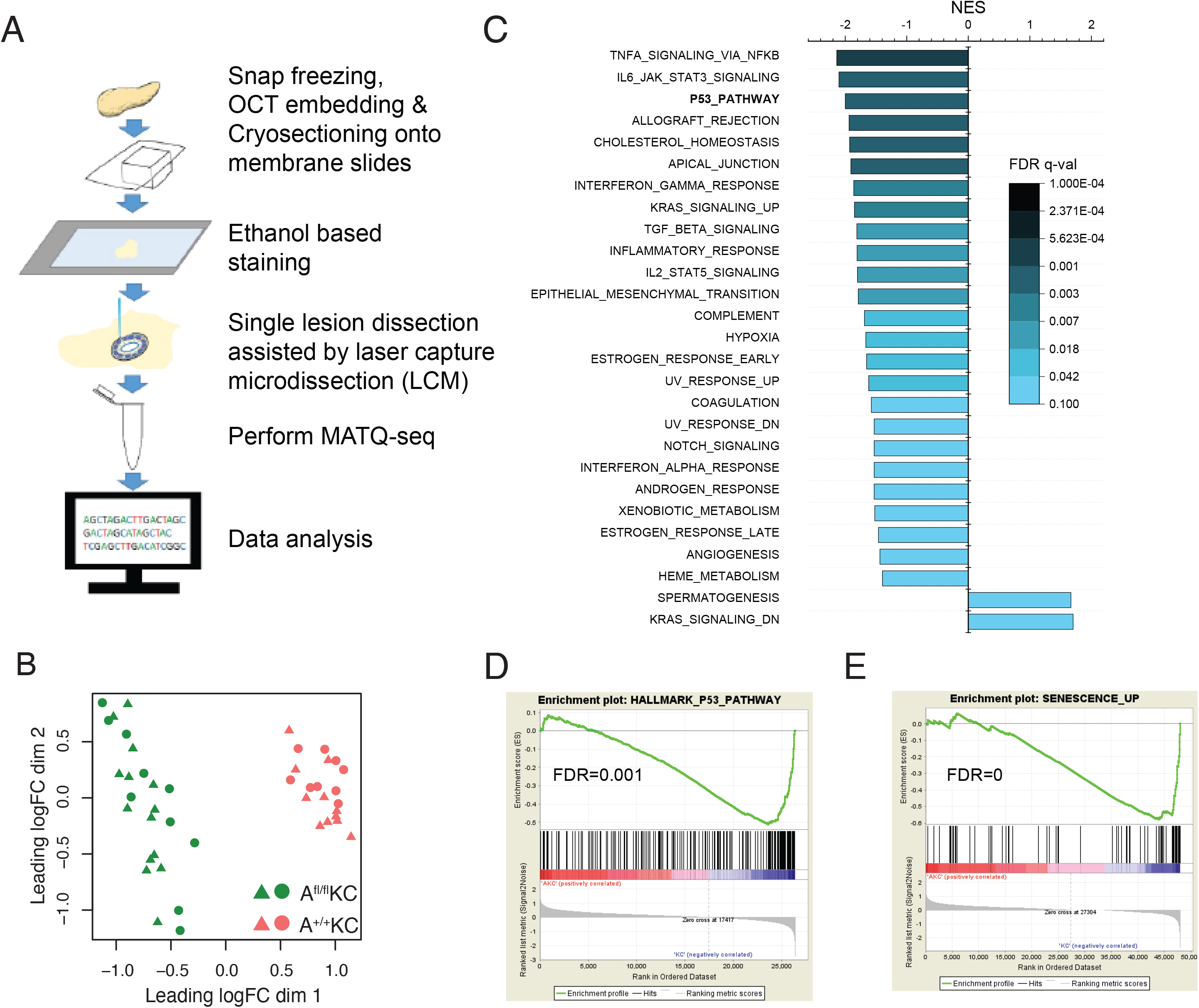
Single PanIN lesion RNA-seq unveils the potential player contributing to the attenuation of Kras-induced senescence in *Arid1a* knockout mice. (**A)** Experimental scheme of transcriptome profiling of single PanIN lesions. (**B**) Multidimensional scaling plot showed a clear separation of the transcriptome profiles of lesions from AKC mice and KC mice. 24 lesions from 2 AKC mice and 20 lesions from 2 KC mice were used for single lesion RNA sequencing. (**C**) Hallmark gene sets that are significantly enriched between lesions from AKC and KC mice. (**D-E**) Enrichment plots of P53_PATHWAY and SENESCENCE_UP from GSEA.

With the differential gene expressions between AKC and KC PanIN lesions, we first performed gene set enrichment analysis (GSEA) using GO biological processes gene sets. As a result, we observed the significant enrichment of genes involved in the development of multiple cellular organisms **(Figure S3A)**. We then interrogated the effects of ARID1A deficiency on the dedifferentiation or trans-differentiation of cells by examining the expression of differentially expressed genes in different mouse tissues (13). As shown in **Figure S3B-C**, the majority of the differentially expressed genes in AKC lesions are highly expressed in the esophagus, liver, colon and also in brain, indicating that a highly complex trans-differentiation occurs following the knockout of Arid1a. Our PanIN-seq approach is capable of characterizing the detailed trans-differentiation at individual PanIN lesion resolution.

With the differentially expressed genes between AKC and KC lesions, next, we performed GSEA using Hallmark gene sets (14) to interrogate the pathways perturbed by *Arid1a* knockout. As a result, we observed that 25 gene sets were downregulated, and surprisingly, only 2 gene sets were upregulated in AKC lesions (**Figure 1C and Figure S4**). Among these changes, it is worth noting that two gene sets are specifically associated with Kras activation: KRAS_SIGNALING_UP and KRAS_SIGNALING_DN.

The gene set KRAS_SIGNALING_UP was downregulated while the gene set KRAS_SIGNALING_DN was upregulated (**Figure S4**). This observation suggests that the activities of Kras signaling are partially suppressed by ARID1A deficiency.

Furthermore, we observed that the p53 signaling pathway is suppressed in AKC lesions (**Figure 1D**). It has been well-established that the upregulation of the p53-related pathway could induce apoptosis or senescence(15). And *ARID1A* mutations have also been shown to be mutually exclusive with TP53 mutations in endometrial cancer (16). Here in order to dissect whether ARID1A is involved in the regulation of apoptosis or senescence, or both, we further examined the activity of related pathways in *Arid1a* KO lesions. Interestingly, only the senescence-associated, but not the apoptosis-associated signaling pathway is significantly suppressed in lesions from AKC mice (**Figure 1E**). This observation led us to hypothesize that ARID1A deficiency could promote the PanIN lesion progression through the attenuation of Kras-induced senescence.

### *In vivo, ex vivo*, and *in vitro* verification of attenuation of Kras-induced senescence by Arid1a deficiency

To verify the effect of ARID1A deficiency on Kras-induced senescence, we first performed senescence-associated beta-galactosidase (SA-β-Gal) staining on the lesions from KC and AKC mice. As a result, SA-β-Gal positive lesions were observed in 5 out of 7 (71%) KC mice. In contrast, only 1 out of 6 (17%) AKC mice showed SA-β-Gal positive lesions. Among the mice with SA-β-Gal positive lesions, the percentage of SA-β-Gal positive lesions in KC mice was nearly two folds of that in AKC mice (**Figure 2A-B)**. These data confirmed that *Arid1a* knockout indeed reduced Kras-induced senescence.

**Figure 2.**
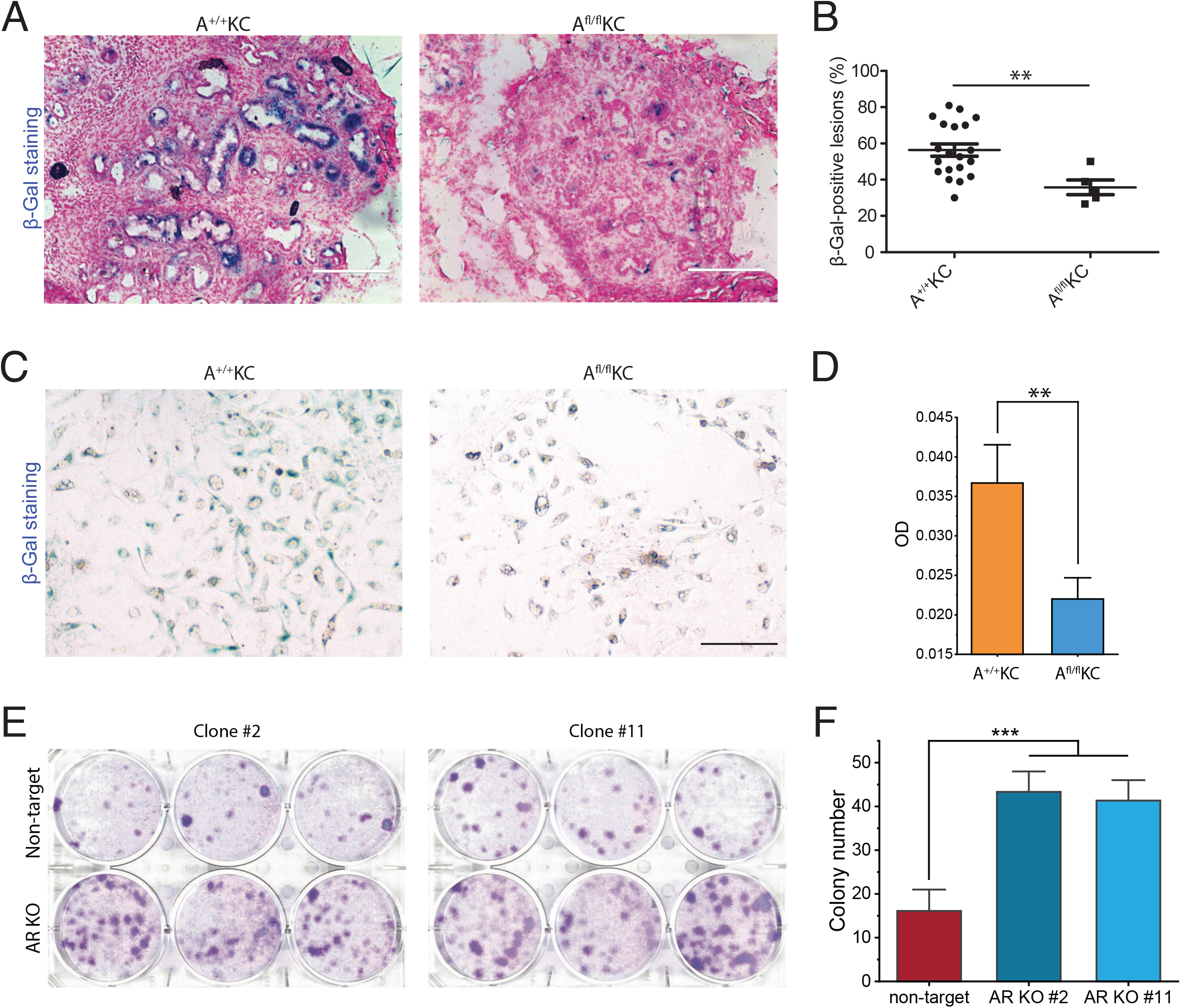
*In vivo, ex vivo,* and *in vitro* verification of attenuation of Kras-induced senescence by *Arid1a* deficiency. (**A**) Representative images of SA-β-gal staining of frozen pancreatic sections from KC mice and AKC mice. (**B**) SA-β-gal positive lesions were counted at 20 random fields under the microscope in four KC mice and 5 random fields from one AKC mouse, presented as a percentage. (**C**) Representative images of SA-β-gal staining of *ex vivo* culture from KC and AKC mice 1 month after administration of tamoxifen. (**D**) Quantification of the intensity of SA-β-gal staining at 10 random fields under the microscope. (**E**) Colony formation assay of *ARID1A* knockout cells and wild type HPNE cells with Kras induction by doxycycline (6ug/ml) for 15 days. (**F**) Quantification of colony number in panel **E**. The colony formation assay was performed twice. Student’s t-test: **, p<0.01; ***, p<0.001. Scale bars, 200 µm.

To further verify the effects of *Arid1a* knockout on senescence, we also performed the *ex vivo* culture experiment of the acinar cells isolated from AKC and KC mice. SA-β-Gal staining was then performed to examine the senescent levels of acinar cells. As shown in **Figure 2C-D**, the intensity of SA-β-Gal staining in the acinar cells from AKC mice was significantly weaker than that from KC mice. More importantly, this result suggests that the effect of *Arid1a* knockout on senescence is likely an intrinsic cellular response of acinar cells, rather than a microenvironmental response such as accelerated clearance of senescent cells by immune cells (17, 18).

Based on the result of the *ex vivo* experiment, next, we tested whether we can observe similar effects of ARID1A deficiency on Kras-induced senescence in *in vitro* cell lines. Here we established a HPNE cell line (human pancreatic Nestin-expressing cells, an intermediary cell type formed during acinar-to-ductal metaplasia) with inducible Kras^G12D^ expression (iKras-HPNE cells) (**Figure S5A**). Next, we knocked out *ARID1A* by CRISPR-Cas9 in iKras-HPNE cells. Two isogenic HPNE clones were used for the following experiments (**Figure S5B**). To examine Kras-induced senescence, we first performed SA-β-Gal staining in *ARID1A* knockout (*ARID1A*-KO) and wild type iKras-HPNE cells. We observed that *ARID1A*-KO cells showed a lower percentage of SA-β-Gal positive cells than the wild type iKras-HPNE cells (**Figure S5C**). The difference is less significant than what we observed both *in vivo* and *ex vivo*. One possible reason for this discrepancy is that in terms of senescence evaluation, SA-β-Gal staining of the immortalized cell lines may not be as effective as in tissue samples (19). Taking an alternative approach, we used the colony formation assay to check the ability of cells to escape from Kras-induced senescence. Indeed, we observed that the knockout of *ARID1A* could drastically increase the number of cells escaping from Kras-induced senescence (**Fig. 2E-F**).

### *ARID1A* knockout significantly upregulates ALDH expression

With the consistent *in vivo, ex vivo*, and *in vitro* observation of anti-senescence effects of ARID1A deficiency, next, we would like to investigate the molecular mechanisms by which *ARID1A* knockout promotes the escape from Kras-induced senescence. We first performed RNA-seq on *ARID1A*-KO (clone #2) and wild type HPNE cells with or without Kras induction. As a result, we observed a clear separation between *ARID1A*-KO and wild type HPNE cells under both conditions in the multidimensional scaling plot (**Figure 3A**). We then analyzed the differentially expressed genes in the *ARID1A*-KO cell line under KRAS induction, and we identified 125 up-regulated genes and 165 down-regulated genes (**Figure 3B & Supplementary Table 2**).

**Figure 3.**
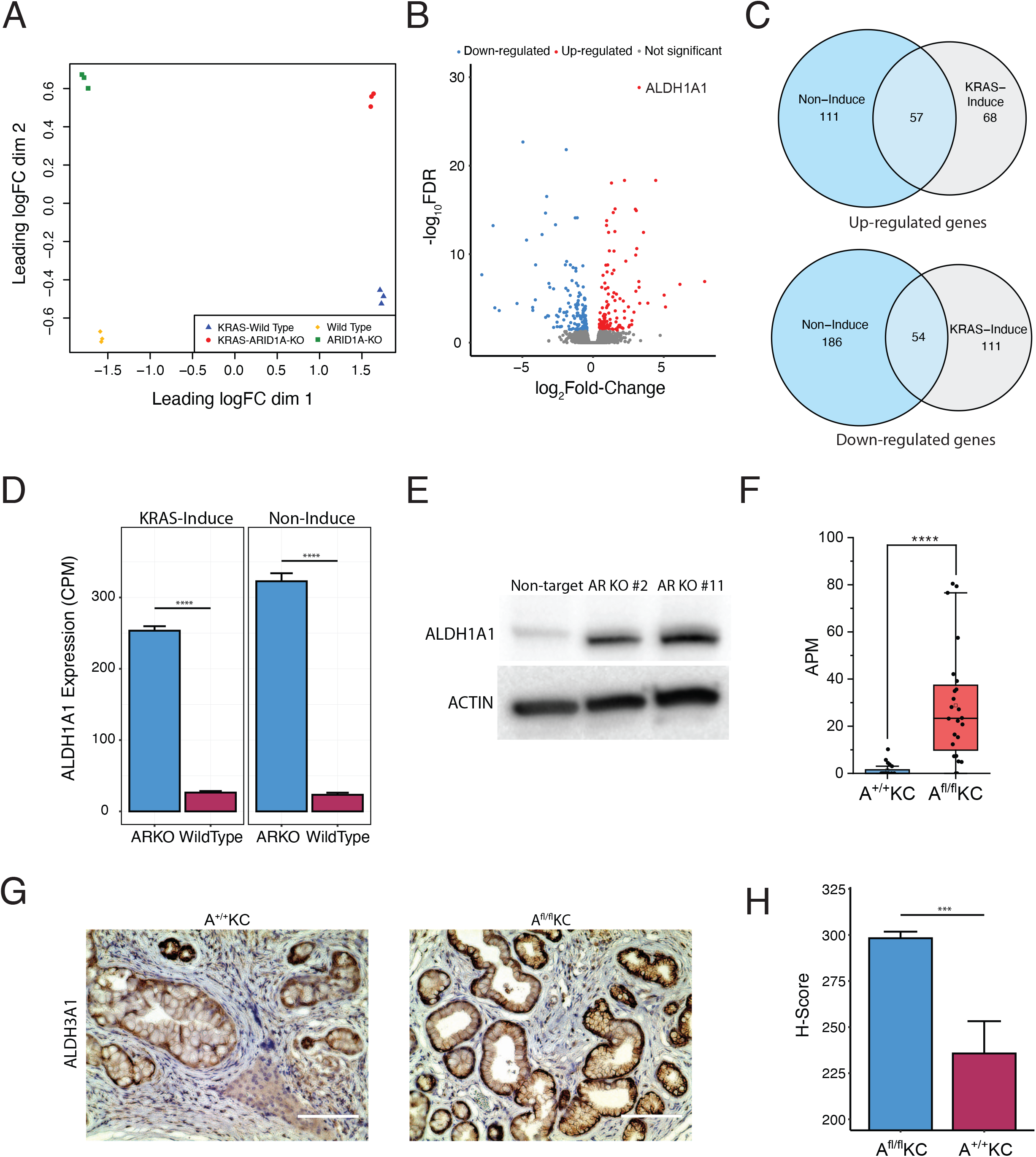
*ARID1A* knockout upregulates ALDH expression. **(A)** Multidimensional scaling plot demonstrated clear separation between the transcriptome profiles of *ARID1A*-KO HPNE cells and the wild type cells with or without Kras induction. RNA sequencing was performed with 3 biological repeats. (**B**) Volcano plot of differentially expressed genes between *ARID1A* knockout cells and wild type cells with Kras induction. (**C**) Venn diagram showing the upregulated genes (upper) and downregulated genes (bottom) that are shared between cells with (grey) or without (blue) Kras induction. (**D**) *ALDH1A1* mRNA levels quantified by sequencing data are significantly different between *ARID1A*-KO cells and wild type cells with (left) or without (right) Kras induction. CPM: count per million reads. (**E**) Western blot for ALDH1A1 expression in *ARID1A*-KO cells and wild type cells with Kras induction. (**F**) mRNA level of *ALDH3A1* in KC and AKC lesions based on PanIN-seq data. APM, amplicon per million reads. (**G**) IHC staining against ALDH3A1 in KC and AKC lesions. Scale bars, 200 µm. (**H**) Comparison of ALDH3A1 levels between KC and AKC lesions based on the intensity of staining in **G**. H-score was calculated by counting the number of lesions with different levels of staining intensity at 10 random fields under the microscope. Student’s t-test: ***, p<0.001; ****, p<0.0001.

To evaluate the gene expression changes that are associated with mutant Kras signaling, we also compared the differentially expressed genes between the cells with or without KRAS induction. As shown in **Figure 3C**, 57 up-regulated genes and 54 down-regulated genes are shared between two conditions. This result shows that about half of upregulated genes and one-third of down-regulated genes are solely dependent on ARID1A deficiency. Therefore, we concluded that there are significant synergistic effects between ARID1A deficiency and mutant KRAS. This observation is also consistent with the PanIN-seq result described above in the partially suppressed Kras signaling pathways due to ARID1A knockout.

Among all the differentially expressed genes between *ARID1A*-KO and wild type HPNE cells, ALDH1A1 has the highest statistical significance with and without KRAS induction (**Figure 3D, Figure S6A, and Supplementary Table 2**). To rule out any potential clonal bias, we also performed RNA-seq on the second clone (clone #11). And we observed that ALDH1A1 was also significantly upregulated in the second clone with and without KRAS induction (**Figure S6B-D and Supplementary Table 2**). The upregulation of *ALDH1A1* in *ARID1A*-KO cells was further verified by both RT-PCR (**Figure S6E**) and western blot (**Figure 3E**).

Next, we examined our PanIN-seq data to evaluate whether similar upregulation can be observed in ALDH expression. Interestingly, we observed that another member of the ALDH family, *Aldh3a1*, is significantly upregulated in lesions from AKC mice (**Figure 3F**). This result was also confirmed by IHC staining (**Figure 3G-H**). It demonstrates a species-dependent difference in the expression of different ALDH proteins.

### ARID1A KO facilitates escape from KRAS-induced senescence via AHDH1A1

Aldehyde dehydrogenase (ALDH) family proteins are known to play major roles in alcohol metabolism by oxidizing aldehydes. However, it is worth noting that over two hundred types of ROS are aldehydes, and many of them are highly toxic (20). Therefore, the ALDH family proteins could also play important roles in reducing these toxic molecules in the cells (21). With the significantly elevated ROS level induced by mutant KRAS, it is possible that ALDH proteins play critical roles in mitigating the cellular oxidative stress. As a result, the highly expressed ALDH proteins can be important for the development of PDAC. Here we analyzed the recent RNA-seq and genomic data of PDAC samples (3) to evaluate whether one of the ALDH family proteins is highly expressed in clinical tumor samples. Indeed, as shown in **Figure 4A**, we observed that in pancreatic cancer, *ALDH1A1* is highly expressed (FPKM > 200) in ~10% of patient samples (81 tumor samples in total). Furthermore, only one patient (0.2% of patient samples, 640 tumor samples from two cohorts (3, 22)) acquired the mutations in *ALDH1A1* (**Figure 4B)**. Both results indicate that ALDH family proteins are not only markers for cancer stem cells but also, they could play important roles in PDAC.

**Figure 4.**
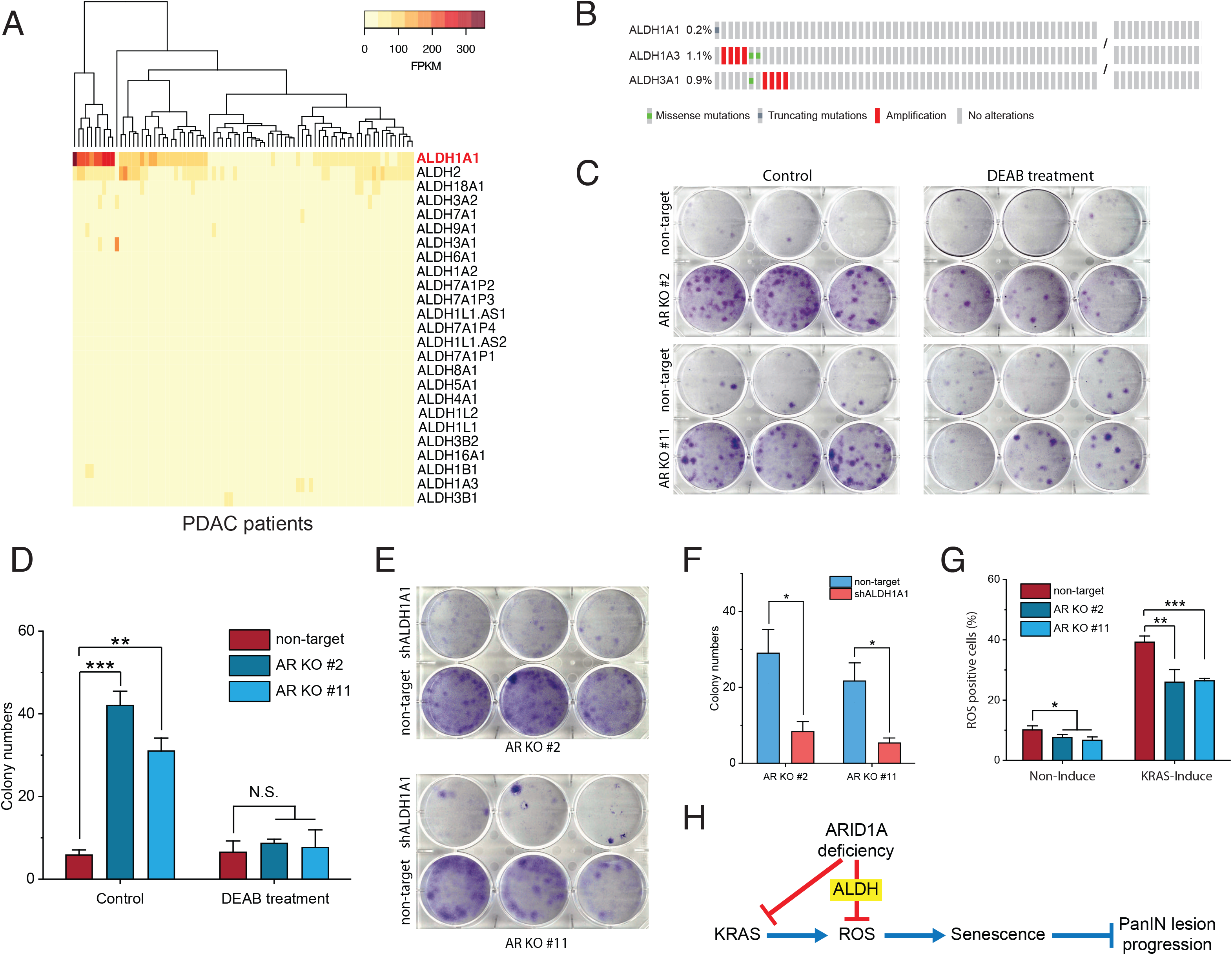
*ARID1A* knockout facilitates escape from Kras-induced senescence by upregulating AHDH1A1 expression. (**A**) Heatmap of the expression levels of ALDH family in PDAC patients (3). (**B**) Mutation rates of ALDH1A1, ALDH1A3, and ALDH3A1 in PDAC patients. The mutation data from these two studies (Bailey et al. (3) and TCGA(4)) are used for analysis. (**C**) colony formation assay of *ARID1A* knockout cells and wild type cells with Kras induction, treated with and without ALDH inhibitor DEAB. (**D**) Quantification of colony number in panel **C**. The colony formation assay was performed twice. (**E**) colony formation assay of *ARID1A* knockout cells expressing shRNA targeting *ALDH1A1* and scramble shRNA control. (**F**) Quantification of colony number in panel **E**. The colony formation assay was performed twice. (**G**) Measurement of the ROS level using a H2DCFDA based ROS detection assay kit. Percentage of positive cells measured by flow cytometry. (**H**) Working model for ARID1A-deficiency-promoted PanIN lesion progression via inhibiting ROS production. Student’s t-test: *, p<0.05; **, p<0.01; ***, p<0.001.

Next, we investigated whether the upregulated ALDH1A1 played an essential role in promoting the escape from Kras-induced senescence. We performed the colony formation assay in HPNE cells with and without N,N-diethylaminobenzaldehyde (DEAB, a pan-inhibitor of ALDH) treatment. As a result, we observed that inhibition of ALDH1A1 activity indeed significantly decreased the number of colonies formed in ARID1A knockout cells; in contrast, no significant changes were observed in the wild type cells (**Figure 4C-D**). To rule out unknown effects of DEAB on HPNE cells, we also performed colony formation assay on *ARID1A*-KO HPNE cells with and without shRNA knockdown of *ALDH1A1*. The knockdown efficiency was verified by RT-PCR (**Figure S6F**). As a result, we also observed that the colony number in *ARID1A*-KO cells with *ALDH1A1* knockdown was significantly less than that without *ALDH1A1* knockdown (**Figure 4E-F**), which is consistent with the result of the ALDH inhibitor experiment.

Furthermore, we examined the levels of ROS production in *ARID1A*-KO cells and wild type cells. And we observed that the fraction of ROS positive cells in *ARID1A*-KO iKras-HPNE cells was significantly less than that in wild type cells regardless of Kras induction (**Figure 4G)**. Supported by both drug inhibition and knockdown experiments, we conclude that *ARID1A* knockout can effectively reduce cellular ROS level by upregulating the expression of ALDH1A1, which then leads to significant attenuation of Kras-induced senescence and acceleration of PanIN progression (**Figure 4H**).

### ARID1A KO activates transcription of the ALDH1A1 gene by increasing the accessibility of its enhancer region

As one of the DNA binding subunits of the SWI/SNF complex, ARID1A can regulate the gene expression by modulation of chromatin structure for transcription factor binding, recruitment of coactivator/corepressor, and facilitation of chromatin looping required to approximate promoters with distal enhancers (6). To investigate how ARID1A deficiency enhances the expression of ALDH1A1, we performed ATAC-seq on *ARID1A*-KO and wild type HPNE cells (**Figure S7 and Supplementary Table 3**). As shown in **Figure 5A**, the Spearman correlation coefficients between knockout and wild type cells are significantly lower than that between replicative samples in both groups, suggesting that the changes of DNA accessibility were robustly captured.

**Figure 5.**
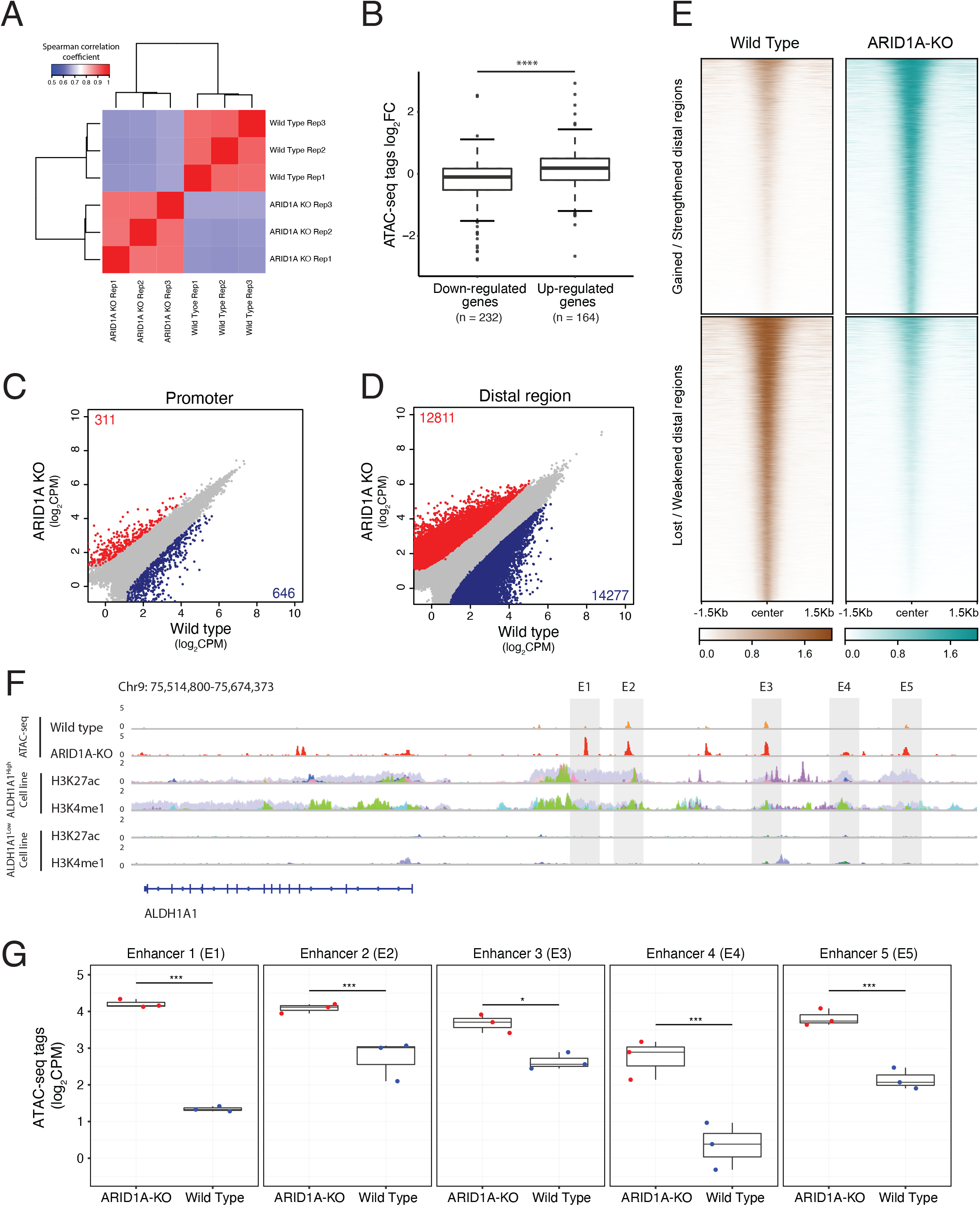
*ARID1A* knockout activates transcription of the ALDH1A1 gene by increasing the accessibility of its enhancer region. (**A**) Spearman correlation coefficients of the read counts in peaks between ARID1A-KO HPNE cells and wild type cells. ATAC sequencing was performed with 3 biological repeats. (**B**) The average fold change of the read counts in the promoters and enhancers of differentially expressed genes identified in RNA-seq. Mann-Whitney-Wilcoxon test: ****, p<0.0001. (**C-D**) The scatter plot of the read counts in each peak between ARID1A-KO cells and wild type cells for promoter (**C**) and distal regions (**D**). The peaks with significantly increased read density in ARID1A-KO cells compared to wild type cells are colored in red. The peaks with significantly decreased read density are colored in blue. (**E**) Heatmap of the gained or lost distal regions between ARID1A-KO cells and wild type cells. (**F**) The ATAC-seq tracks and H3K4me1/H3K27ac ChIP-seq tracks of the distal regions of the *ALDH1A1* gene. The ChIP-seq tracks from different cell lines are labeled in different colors and overlaid. The figure with separated tracks is shown in **Supplementary Figure 9**. The ALDH1A1^high^ cell lines include A549, LOUCY, A673, 22RV1, VCAP, K562, and HepG2. The ALDH1A1^low^ cell lines include HCT116, MCF7, Panc1, PC9, and DOHH2. The ChIP-seq data were obtained from the ENCODE database (33). (**G**) Read counts in the 5 enhancer peaks of the *ALDH1A1* gene in ARID1A-KO HPNE cells and wild type cells. P-value: *, p<0.05; ***, p<0.001; ****, p<0.0001.

We first examined the accessibility of the promoter and enhancer regions of the genes that are differentially expressed between the *ARID1A*-KO cells and the wildtype cells. And we observed that the changes in the accessibility of regulatory regions were significantly correlated with the alterations in gene expression levels in ARID1A-KO cells (**Figure 5B**). This result buttresses that ARID1A knockout alters the gene expression by modulating the chromatin accessibility of the regulatory elements. Next, when we analyzed the regulatory regions with significant changes in accessibility between ARID1A-KO and wild type cells, we observed that ARID1A deficiency mainly affects the chromatin accessibility at distal regulatory regions (**Figure 5C-E and Figure S8**). The significant interactions between SWI/SNF complex and distal regulatory regions such as enhancers have also been observed in colon cancer (23).

Next, we examined the accessibility of the promoter and distal regulatory elements of the ALDH1A1 gene in both ARID1A-KO and wild type cells. Interestingly, we found that there is a significant increase in the accessibility in the 9 out of 11 peaks at the distal regions when we compared ARID1A-KO cells to wildtype cells (**Supplementary Table 4**). To further verify these functional regions, we also compared the landscape of two well-known enhancer markers (H3K27ac and H3K4me1) in seven ALDH1A1 highly expressed cell lines with our ATAC-seq peaks (**Supplementary Table 5**). As a result, we observed that five loci are clearly overlapped with the active enhancer markers (**Figure 5F and Figure S9**). As a comparison, we also plotted out the landscape of H3K27ac and H3K4me1 in five ALDH1A1 lowly expressed cell lines, and we did not observe any significant overlap between our ATAC-seq peaks and H3K27ac/H3K4me1 enriched regions (**Figure 5F**). Furthermore, we also observed the consistent increase of accessibility for the 5 enhancer loci in ARID1A knockout cells in comparison to the wildtype cells (**Figure 5G**). Overall, these results confirm that ARID1A deficiency upregulates ALDH1A1 expression by increasing the accessibility of the associated enhancer elements.

To identify the proteins that could potentially bind to these 5 enhance loci, we examined the transcription factor (TF) ChIP-seq datasets from 7 ALDH1A1^high^ cell lines and 5 ALDH1A1^low^ cell lines (24). We counted the binding events of each TF in the five candidate enhancer peaks for both ALDH1A1^high^ cell lines and ALDH1A1^low^ cell lines. And we list the TFs that could bind to the enhancer regions in **Figure S10**. For the TFs whose binding events are preferentially detected in the datasets from ALDH1A1^high^ cell lines (Fisher’s test, p < 0.05), we marked them in red color, which includes EP300 and NR3C1. For the TFs that do not have enough datasets for a statistical test, we marked the TFs whose binding events are observed in more than 50% of ALDH1A1^high^ cell line sets in blue color; and the TFs whose binding events are observed in more than 25% but less than 50% of ALDH1A1^high^ cell line sets in black color. The TFs that are not expressed in HPNE cell lines were excluded.

## DISCUSSION

With KRAS as a *bona fide* driver oncogene in pancreatic cancer, how to counteracts the oncogenic stress induced by mutant KRAS is critically important for the progression of precursor lesions (25). It has been shown that Kras itself can upregulate the expression of multiple oxidoreductases via Nrf2 (nuclear factor, erythroid derived 2, like 2) to counteract the oncogenic stress (26). Meanwhile, Kras can also enhance NADPH production by reprogramming the metabolism of glutamine to assist oxidoreductases in scavenging ROS (27). However, the observation that a high percentage of senescent PanIN lesions occur in the pancreas from KC mice suggests the mechanisms described above likely have limited effects in reducing Kras-induced ROS in neoplastic lesions. In the early development of PDAC, it is plausible that new mutations can be acquired to help cells to reduce ROS and escape the oncogenic KRAS induced senescence.

In this study, we successfully adapt a high-sensitivity total-RNA-based single-cell RNA-seq method: MATQ-seq to profile the transcriptome of single lesions. Supported by the total RNA-detection ability of MATQ-seq (12), we achieved high sensitivity in gene detection for individual early-stage lesions with 10-30 cells per lesion despite the RNA degradation frequently occurred in pancreatic samples. The transcriptome profiling of the early-stage PanIN lesions allows us to dissect the signaling pathways involved in PanIN progression directly. And we found that *ARID1A* knockout can significantly reduce the levels of senescence. Using the HPNE cell line model, we further showed that the upregulation of ALDH proteins plays an essential role in reducing ROS levels and promoting the attenuation of senescence.

It is worth noting that *ARID1A* has also been linked to the regulation of ROS through other pathways. Sun *et al*. found ARID1A overexpression causes an increase of ROS by activating transcription of cytochrome P450 enzymes (CYP450) at the initiation stage of liver cancer (28). Interestingly, Ogiwara *et al.* found ARID1A deficiency results in elevated ROS by inhibiting the transcription of *SLC7A11* (a transporter gene required for the import of cystine and the production of glutathione) in ovarian cancer (29). Therefore, the effects of *ARID1A* deficiency on ROS is likely tissue specific.

Besides the reduced ROS levels, we also observed that ARID1A deficiency suppressed the activation of several inflammation-or immune-related gene sets in the PanIN lesion. As one of the main phenotypes of senescent cells, senescence-associated secretory phenotype (SASP) involves secretion of various cytokines, especially proinflammatory cytokines (30). As a result, with the attenuation of Kras-induced senescence, we would expect to observe reduced inflammatory responses due to ARID1A deficiency. Indeed, in the RNA-seq data of both PanIN lesions and HPNE cells with ARID1A deficiency, we observed the downregulation of NF-kB signaling (the master regulator of SASP maintenance (31)).

Beyond counteracting the ROS levels induced by oncogenic KRAS, the upregulated expression of ALDH proteins could also reduce the efficacy of drug treatment since many chemotherapy drugs promote cell death by inducing ROS and DNA damage in tumor cells (32). For patients with Arid1a mutations or deletion, or high-level expression of ALDH proteins, ALDH-mediated ROS scavenging may significantly impair the efficacy of these drugs and lead to the drug-resistance. Therefore, it is greatly desired to study the efficacy of the combination of ALDH inhibitors and chemotherapy drugs in reducing drug resistance and tumor relapse in the future.

Overall, in this study, we showed that transcriptome profiling of individual lesions is not only technically feasible, but also it can provide important insights about the potential mechanisms in tumor progression. Our results unveil that *ARID1A* knockout can significantly reduce Kras-induced senescence by upregulating the expression of ALDH family proteins. We showed that ALDH family proteins are not only just marker proteins for cancer stem cells; more importantly, they play essential roles in mitigating the ROS stress induced by oncogenic KRAS.

## ACKNOWLEDGEMENTS

We are grateful to the McNair family and Dr. C. Neblett for their support. The development of PanIN-seq was supported by the NIH Director’s New Innovator Award (1DP2EB020399). We thank Dr. Zhong Wang (University of Michigan) for providing the Arid1a floxed mice. We thank Kuanwei Sheng, Ejune Chen, Jinxiang Yuan, and other Zong lab members for their help in this project. We thank Dr. Christophe Herman for his proofreading and helpful discussion.

## CONTRIBUTIONS

C.Z. designed the project. S.L. W.C., Y.N., and C.Z. wrote the manuscript. S.L., W.C., J.L., Y.Z., and Z.H. contributed to the mice model development. W.C. and Y.N. performed single-PanIN-seq experiments. S.L. performed *ex vivo* and *in vitro* experiments. Y.N. performed ATAC-seq. W.C. and Y.N. performed the sequencing analysis.

