## Supplemental Materials for "Single-PanIN-seq Unveils that ARID1A Deficiency Promotes Pancreatic Tumorigenesis by Attenuating KRAS Induced Senescence"

**Of**

### MATERIAL AND METHODS

#### Mice

All animal experiments in this study were performed following a protocol approved by IACUC of Baylor College of Medicine. In this study, the following mice strains were generated: *Arid1a*$^{fl/fl}$*;LSL-Kras*$^{G12D/+}$*;Ptf1a*$^{CreERT/+}$, *Arid1a*$^{fl/+}$*;LSL-Kras*$^{G12D/+}$*;Ptf1a*$^{CreERT/+}$, and *Arid1a*$^{+/+}$*;LSL-Kras*$^{G12D/+}$*;Ptf1a*$^{CreERT/+}$. The promoter of *Ptf1a* is used to drive the expression of inducible Cre in adult acinar cells (*Ptf1a*$^{CreERT/+}$, denoted as *C*). Removal of the floxed exon 8 of the *Arid1a* (denoted as *A*$^{fl}$) leads to loss of *Arid1a* transcript (29). Oncogenic Kras mutant allele is silenced by a floxed STOP cassette (*LSL-Kras*$^{G12D/+}$, denoted as *K*). The adult mice (6-8 weeks) of the above genotypes were administered with tamoxifen (75 mg/kg/d for 5 consecutive days) to induce efficient ablation of ARID1A and activation of oncogenic Kras in pancreatic acinar cells.

#### Cell lines

HPNE was purchased from ATCC and cultured in media recommended by ATCC. HPNE with inducible Kras$^{G12D/+}$ (HPNE-Kras) was generated by the transfection of plasmid pInducer20-Kras$^{G12D}$ into parental HPNE cells. After G418 selection for 15 days, the survived cells were then used for single-cell expansion in a 96-well plate. For *ARID1A* knockout HPNE cells, an isogenic clone of HPNE-Kras was infected with lentivirus packaged with pL-CRISPR.EFS.tRFP-ARID1A. After infection for 5 days, the RFP-positive cells were sorted, and single-cell expansion was performed. *ARID1A* knockout was confirmed by Sanger sequencing of the isogenic clones. The guide RNA sequence used for *ARID1A* knockout is CAGCGGTACCCGATGACCAT. For *ALDH1A1* knockdown cells, two *ARID1A* knockout HPNE clones were infected by lentivirus packaged with plasmid pGIPz-*ALDH1A1*. After infection for 5 days, the GFP-positive cells were sorted. Knockdown efficiency was confirmed by RT-PCR. The shRNA sequence targeting *ALDH1A1* is GGAGTGTTTACCAAAGACATT.

#### Primary acinar isolation and culture

Pancreata isolated from KC mice and AKC mice 1 month after 5-day tamoxifen administration were rinsed twice in cold 1×HBSS buffer, then minced into small pieces and digested with digestion buffer (HBSS buffer with 10 mM HEPES and 0.5 mg/ml collagenase and 0.25 mg/ml trypsin inhibitor) for 20-30 minutes at 37 °C. During the incubation. The tissue was pipetted every 5 minutes. After washing twice with washing buffer (HBSS buffer with 5% FBS), the digested tissue was resuspended with media (Waymouth media with 2.5% FBS and 0.25 mg/ml trypsin inhibitor and 100 U/mL Penicillin-Streptomycin), filtrated with 100 $\mu$m strainer and seeded into 10cm dishes overnight at 37 °C to remove fibroblasts and ductal cells. The unattached acinar cells were then transferred into collagen-coated plates for growth. To activate Kras expression, 25 ng/ml EGF was added into the media for 5 days. Cells were then used for SA-β-Gal staining, RT-PCR, and western blot.

**Western blot**

Western blot was performed using the standard protocol. Antibodies used in this study include ALDH1A1 (Abcam, Cat.# ab23375), ALDH3A1 (Abcam, Cat.# ab76976), Kras (Abcam, Cat.# ab180772), phosph-Erk1/2 (Cell Signaling Technology, Cat.# 4370S), ARID1A (Santa Cruz Biotechnology, Cat.# sc-32761) and β-actin (Sigma-Aldrich, Cat.# A1978).

**Colony formation assay**

HPNE cells ($3\times10^4$/well) were seeded into 6-well plates. The cells were treated with doxycycline (6$\mu$g/ml for 15 days) with and without DEAB (1.5 $\mu$M for 30 days). The media was changed every 2 days. When the colonies were large enough, the cells were fixed and stained with crystal violet.

**ROS measurement**

HPNE cells were treated with doxycycline (6$\mu$g/ml for 5 days). ROS level was measured by Flow Cytometry using ROS Detection Assay Kit (BioVision, Cat.# K936-250).

**SA-β-Gal staining**

SA-β-Gal staining was performed on slides of freshly frozen tissues or cells using Senescence β-Galactosidase Staining Kit (Cell Signaling Technology, Cat.# 9860). Total and SA-β-Gal positive lesions or cells were counted at random fields under the microscope, and positive rates were calculated. For quantification of SA-β-Gal staining of primary acinar cells, due to the difficulty of recognizing the nuclei, average optical density (OD) was used to quantify the intensity of SA-β-Gal staining. 8-bit images were adjusted for white balance and color-deconvoluted using the Feulgen light green vector in ImageJ. The average grey values of the green channel were measured. Optical density was calculated using the following formula: OD=log10 (255/grey value).

**IHC quantification**

For the quantification of IHC results against ALDH3A1, the H-score method was used. In brief, staining intensity (not stained: 0, weakly stained: +1, moderately stained: +2, or strongly stained: +3) was determined for each lesion of interest in the field. The H-score was calculated by the following formula: 3 x percentage of strongly stained cells + 2 x percentage of moderately stained cells + 1 x weakly stained cells, giving a range of 0 to 300.

**Bulk RNA-seq**

HPNE cells were treated with doxycycline (6$\mu$g/ml) for 5 days. RNA samples were prepared using the standard protocol for Trizol. mRNA was enriched using NEBNext Poly(A) mRNA Magnetic Isolation Module (NEB, E7490), and the library was prepared using the NEBNext Ultra II RNA Library Prep Kit for Illumina (NEB, E7770). All libraries were sequenced on Illumina Nextseq500 platform. Reads were aligned to hg19 assembly of the human genome by STAR aligner (30), and transcripts counting was performed by HTseq-count (31). Differential gene expression analysis was performed by using edgeR (32) with a cutoff of FDR at 0.05.

### ATAC-seq experiment

ATAC-seq was performed following the protocol of Howard Chang's lab (https://www.nature.com/articles/nmeth.4396) with slight modifications. In brief, $5x10^4$ cells were lysed with ATAC-Resuspension Buffer (RSB) containing 0.1% NP40 and 0.1% Tween-20. After incubation on ice for 3 minutes, the cell lysates were washed by RSB with 0.1% Tween-20. The cell lysates were then incubated with transposition mixture at 37°C for 30 minutes. After amplification, the transposed fragments were purified with magnetic beads. Finally, 4ng fragments were used for the generation of the library. All libraries were sequenced on Illumina Nextseq500 platform.

### ATAC-seq data analysis

Reads were then mapped to the hg19 assembly by Bowtie2 (33) after removing the adaptor sequence. The quality control of ATAC-seq data was performed by using the ATACseqQC R package (34). Next, the mapped reads from three technical replicates of each genotype were combined for the peak calling by MACS2 (35). Peaks from wild type samples and *ARID1A-KO* samples were combined to get a union peak set. All the peaks were then annotated by HOMER (36). HTseq-count (31) was used for read counting. edgeR package (32) was then used for normalization between different samples and for peak differential analysis. The read density profiles of the differential peaks were plotted by deepTools (37).

### ENCODE data analysis

The expression levels of *ALDH1A1* in the cell lines in ENCODE were obtained from the Cancer Cell Line Encyclopedia (CCLE) database (38). Seven cell lines with RPKM > 10 were categorized as $ALDH1A1^{high}$ cell lines. Five cell lines with RPKM < 0.5 were selected as $ALDH1A1^{low}$ cell lines. The A549 H3K27ac and H3K4me1 ChIP-seq data were reanalyzed by using hg19 assembly as described above. The peak files of H3K27ac/H3K4me1 from the other cell lines were directly obtained from the ENCODE database. The ALDH1A1-related differential ATAC peaks, which are overlapped with the

H3K27ac/H3K4me1 peaks in at least 4 out of 7 ALDH1A1$^{high}$ cell lines, were characterized as functional enhancer regions.

**Laser capture microdissection and MATQ-seq of single lesions**

We used the MMI CellCut platform to perform Laser capture microdissection **(**LCM). 40% to 50% laser power was used with the cutting speed of 18 $\mu$m/s to dissect microscopic lesions. 1.6 $\mu$L of MATQ-seq lysis buffer was added onto the isolation cap where the dissected tissue was attached (MMI, Prod. No. 50206). We used a pipette tip to scrape the laser dissected tissue into the lysis buffer and then pipetted the lysis buffer into the tube. Sample tubes were then placed on a thermocycler and incubated at 72°C for 3.5 minutes, followed by 1 min incubation on ice. 2.4 $\mu$L of MATQ-seq RT buffer was then added. MATQ-seq and standard library prep were performed. All libraries were sequenced on Illumina Nextseq500 platform.

**MATQ-seq data analysis**

The raw sequencing data trimming and barcode retrieval were performed as previously described (12). The reads were mapped to the genome MM10 using STAR with the following parameters: --outFilterMismatchNoverLmax 0.05 --outFilterMatchNmin 16 --outFilterScoreMinOverLread 0 --outFilterMatchNminOverLread 0. We used Gencode annotation release M10 (GRCm38.p4) for transcript annotation. Unique barcode counting and gene expression level quantification were performed as previously described with a few modifications: the mapping position of the reads was included as part of the identity of the corresponding barcodes; only reads mapped to the exon region were used for gene expression level quantification. Differential gene expression analysis was performed by using edgeR with a cutoff of FDR at 0.05.

For mouse tissue expression pattern analysis, we used data sets from Söllner et al. (13). THE average TPM for each tissue was calculated and log transformed for the later analysis. Hierarchical clustering was performed using the Clustergram function in

MATLAB. Standardization was performed for each gene, and Euclidean distance was used for clustering.

#### Functional enrichment analysis

Functional enrichment analysis was performed by using GSEA with the MSigDB hallmark gene sets, and the senescence-related gene sets from the MSigDB curated gene sets.

#### Sequencing Data

Individual PanIN lesion RNA-seq data and bulk RNA-seq data are deposited in GEO under GSE160444.

To review GEO accession GSE160444:
Go to https://urldefense.proofpoint.com/v2/url?u=https-3A__www.ncbi.nlm.nih.gov_geo_query_acc.cgi-3Facc-3DGSE160444&d=DwIBAg&c=ZQs-KZ8oxEw0p81sqgiaRA&r=s2BDbzIQ19J3W-_I3taw4ILuEShXAGQ5Zy_WqXCcGdQ&m=O1p04Be7qzzSiKQ25cvlGHMVzW4q1g8kpVSo0HbN6G8&s=8Qf6FybrhH-1M9rM0QQFjSNY4tU2-3ahSkpwGDzQ4eo&e=
Enter token ytwbmoqcndavroz into the box

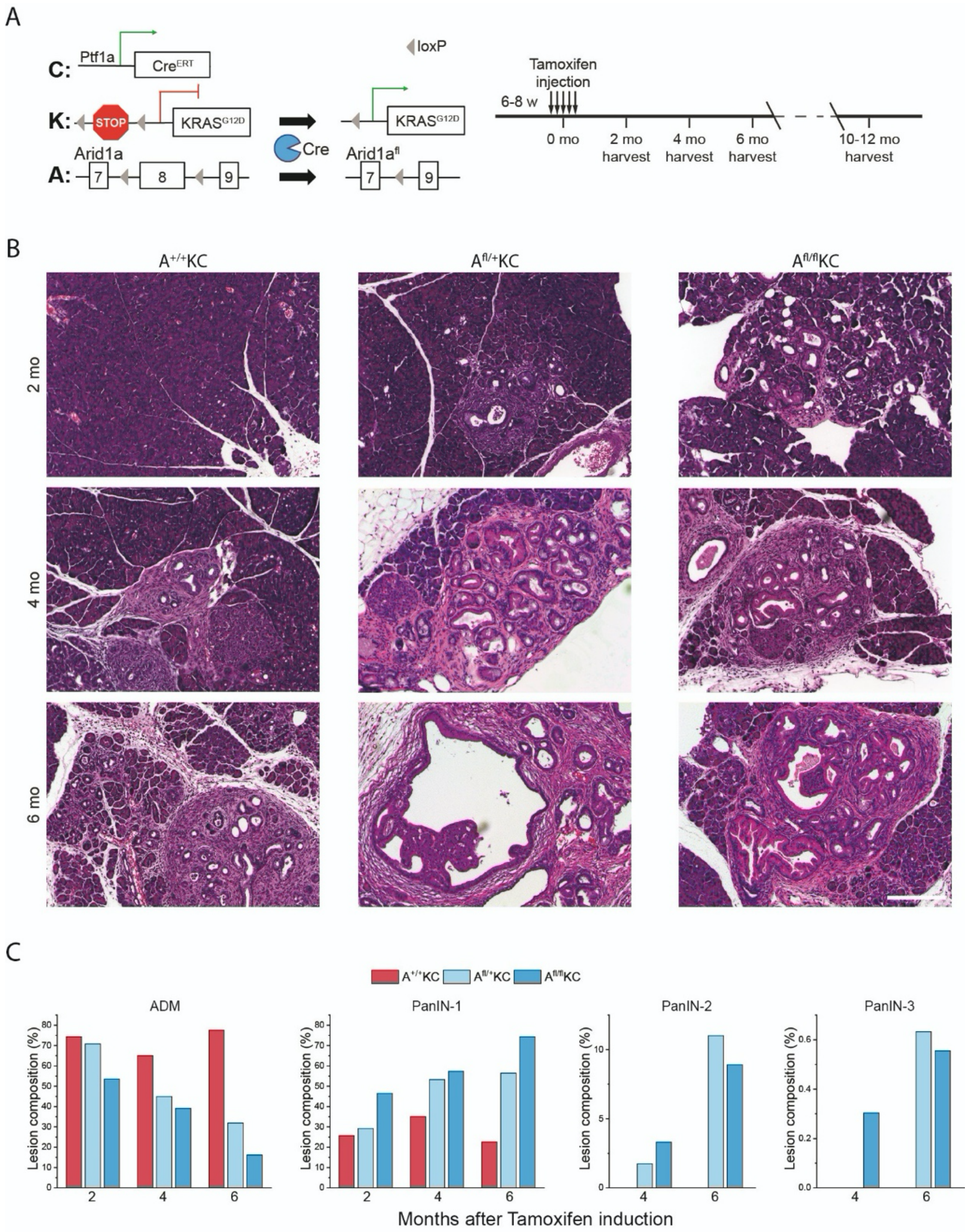


**Figure S1.** *Arid1a* knockout drastically accelerates PanIN progression. (**A**) Genetic makeup and experimental scheme. (**B**) Representative H&E staining images of pancreata from $A^{+/+}$KC, $A^{fl/+}$KC, and $A^{fl/fl}$KC mice after administration of tamoxifen for 2, 4, and 6 months. (**C**) Lesions with different grades were counted at 10 random fields under the microscope, presented as a percentage. Scale bar, 200 $\mu$m.

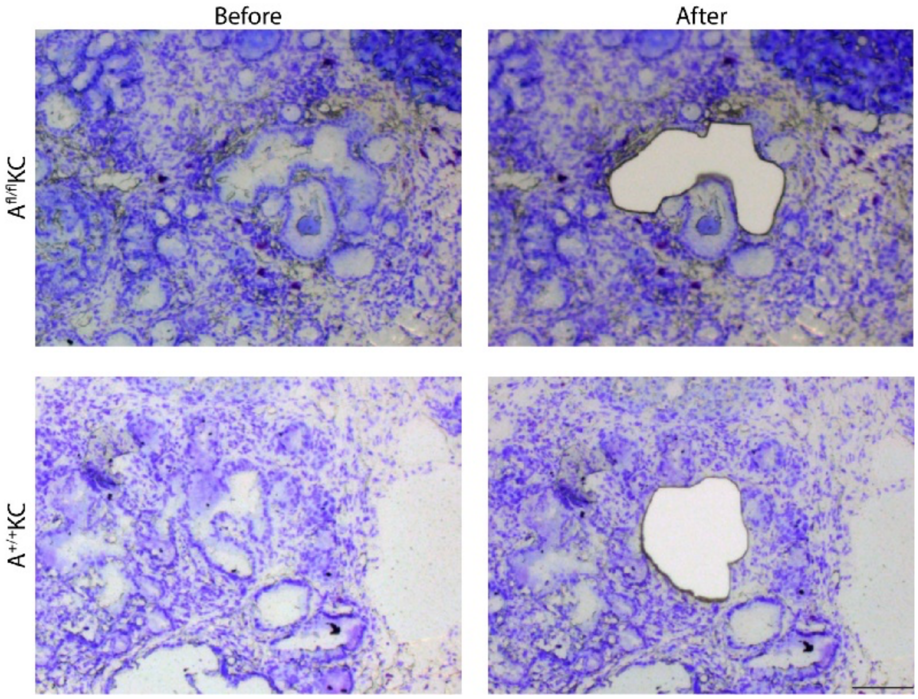


**Figure S2.** Representative images of the lesion region before and after laser capture microdissection. Scale bar, 100 $\mu$m.

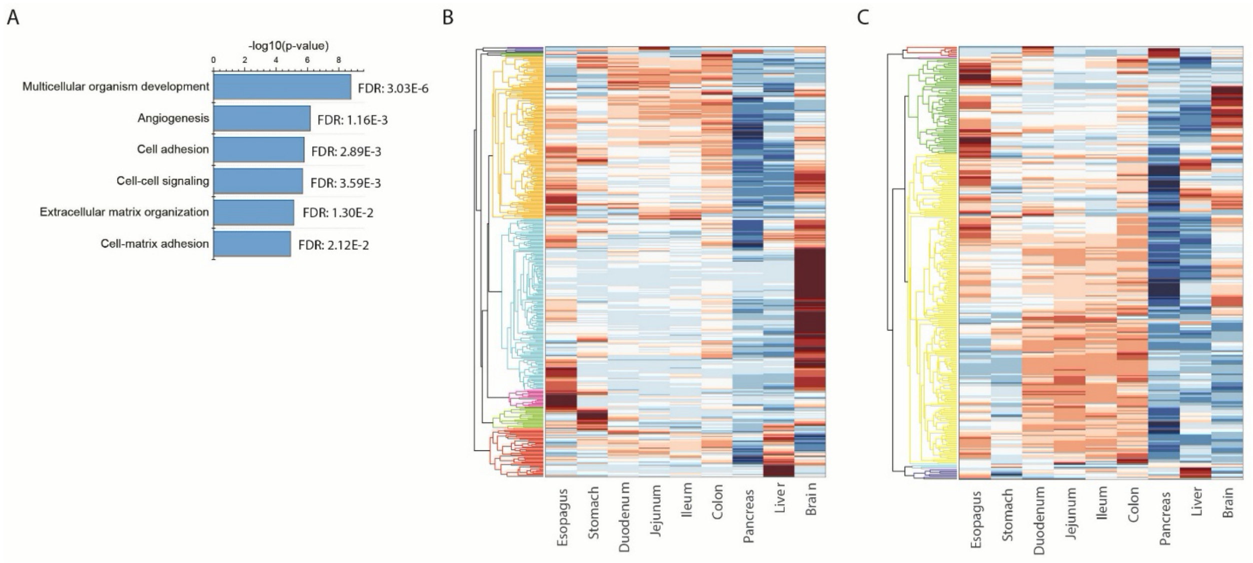


**Figure S3.** (**A**) Functional enrichment analysis of up-regulated genes in $A^{fl/fl}KC$ lesions in GO: biological processes performed by DAVID. **(B-C)** Mouse tissue expression pattern of the up- and down-regulated genes in $A^{fl/fl}KC$ lesions (data source: Söllner *et al*. Scientific Data, 2017).

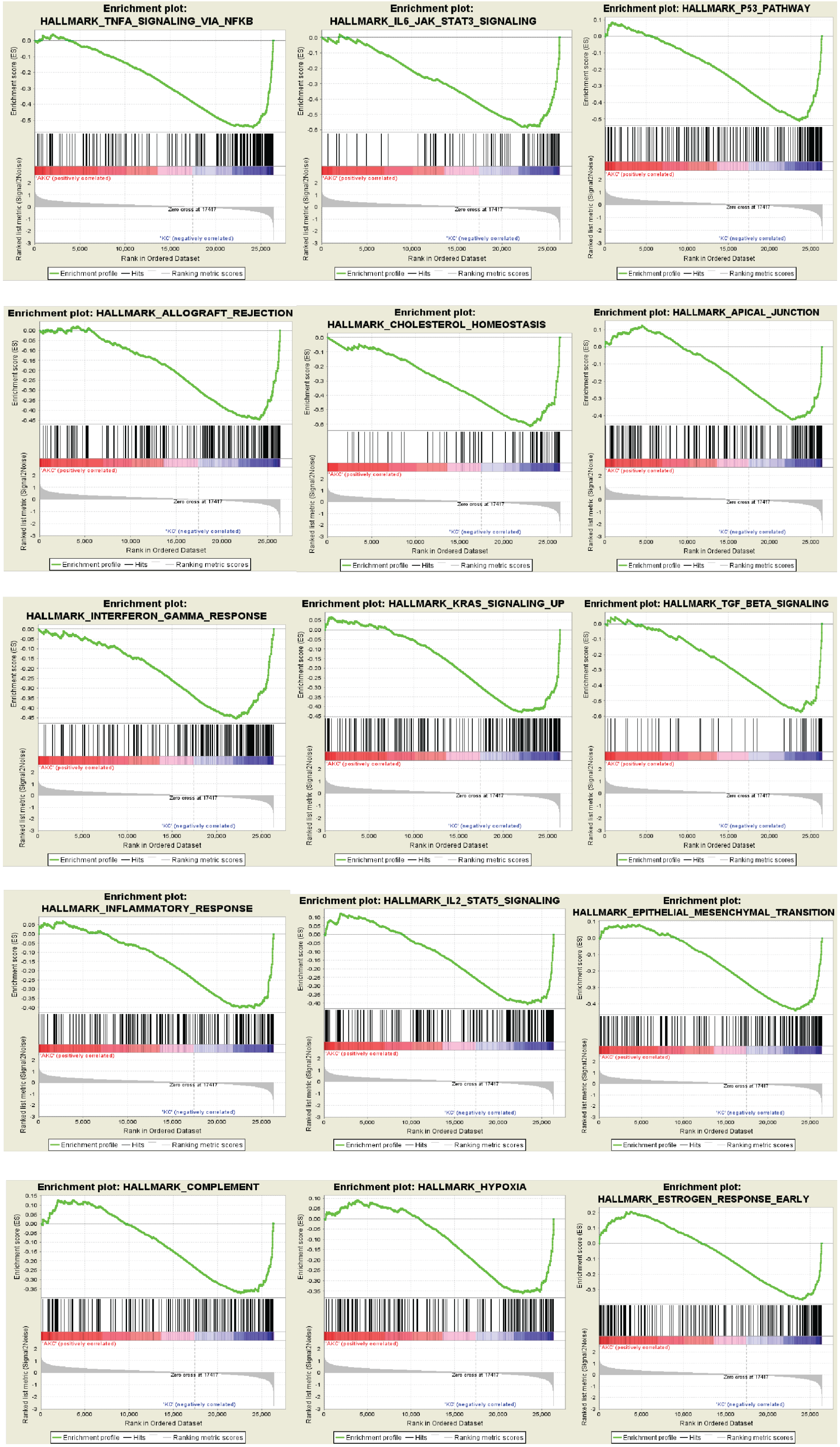
Enrichment plot: HALLMARK_TNFA_SIGNALING_VIA_NFKB
Enrichment plot: HALLMARK_IL6_JAK_STAT3_SIGNALING
Enrichment plot: HALLMARK_P53_PATHWAY
Enrichment plot: HALLMARK_ALLOGRAFT_REJECTION
Enrichment plot: HALLMARK_CHOLESTEROL_HOMEOSTASIS
Enrichment plot: HALLMARK_APICAL_JUNCTION
Enrichment plot: HALLMARK_INTERFERON_GAMMA_RESPONSE
Enrichment plot: HALLMARK_KRAS_SIGNALING_UP
Enrichment plot: HALLMARK_TGF_BETA_SIGNALING
Enrichment plot: HALLMARK_INFLAMMATORY_RESPONSE
Enrichment plot: HALLMARK_IL2_STAT5_SIGNALING
Enrichment plot: HALLMARK_EPITHELIAL_MESENCHYMAL_TRANSITION
Enrichment plot: HALLMARK_COMPLEMENT
Enrichment plot: HALLMARK_HYPOXIA
Enrichment plot: HALLMARK_ESTROGEN_RESPONSE_EARLY
Enrichment score (ES)
Ranked list metric (Signal2Noise)
'AKC' (positively correlated)
Zero cross at 17417
'KC' (negatively correlated)
Rank in Ordered Dataset
Enrichment profile
Hits
Ranking metric scores

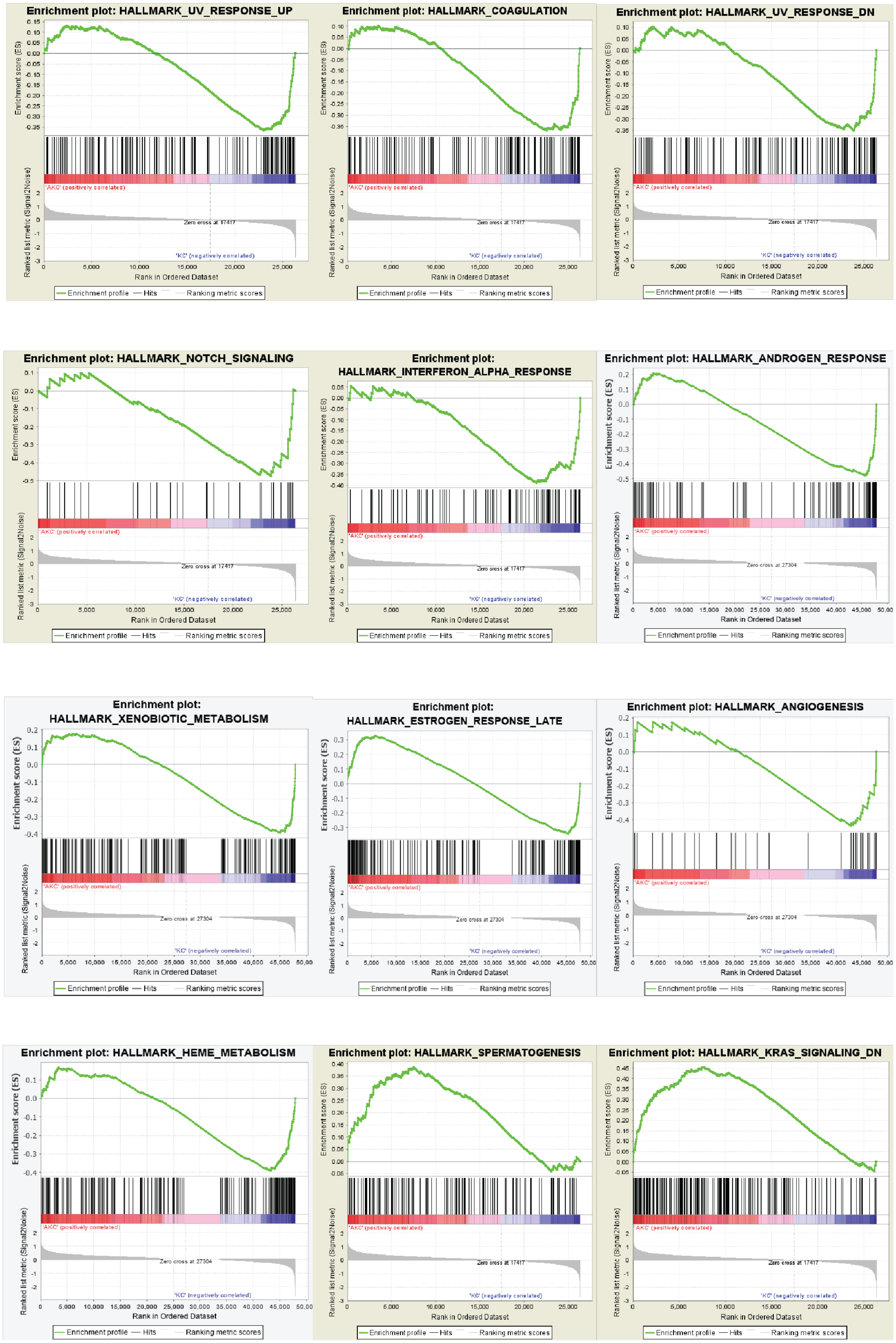
Enrichment plot: HALLMARK_UV_RESPONSE_UP
Enrichment plot: HALLMARK_COAGULATION
Enrichment plot: HALLMARK_UV_RESPONSE_DN
Enrichment plot: HALLMARK_NOTCH_SIGNALING
Enrichment plot: HALLMARK_INTERFERON_ALPHA_RESPONSE
Enrichment plot: HALLMARK_ANDROGEN_RESPONSE
Enrichment plot: HALLMARK_XENOBIOTIC_METABOLISM
Enrichment plot: HALLMARK_ESTROGEN_RESPONSE_LATE
Enrichment plot: HALLMARK_ANGIOGENESIS
Enrichment plot: HALLMARK_HEME_METABOLISM
Enrichment plot: HALLMARK_SPERMATOGENESIS
Enrichment plot: HALLMARK_KRAS_SIGNALING_DN
Enrichment score (ES)
Ranked list metric (Signal2Noise)
'AKC' (positively correlated)
'KC' (negatively correlated)
Zero cross at 17417
Zero cross at 27304
Rank in Ordered Dataset
Enrichment profile
Hits
Ranking metric scores

**Figure S4.** The genes enrichment plots of 27 relevant pathways with statistical significance in A$^{fl/fl}$KC lesions.

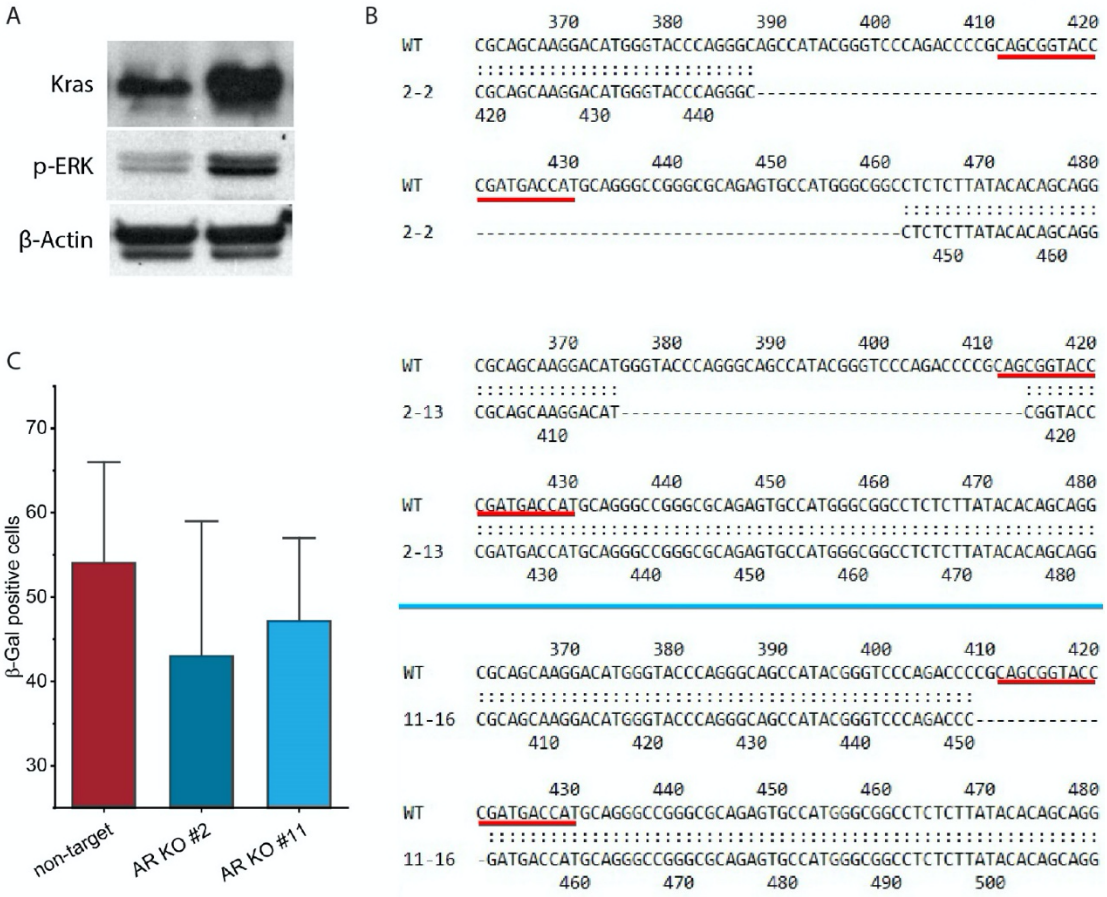


**Figure S5.** Generation of the cell line with inducible Kras overexpression and ARID1A knockout. (**A**) Examination of activation of Kras signaling by western in HPNE cells with inducible Kras knockin (iKras-HPNE cells clone #4). Phosphorylation of ERK was used as the indicator for Kras activation. (**B**) Confirmation of *ARID1A*-KO by Sanger Sequencing in iKras-HPNE cells clone #4 plus *ARID1A*-KO (clone #2 and #11). The underscored sequence is the target for guide RNA. (**C**) SA-βGal staining in HPNE cells with *ARID1A*-KO (clone #2 and #11) and non-target control upon Kras induction.

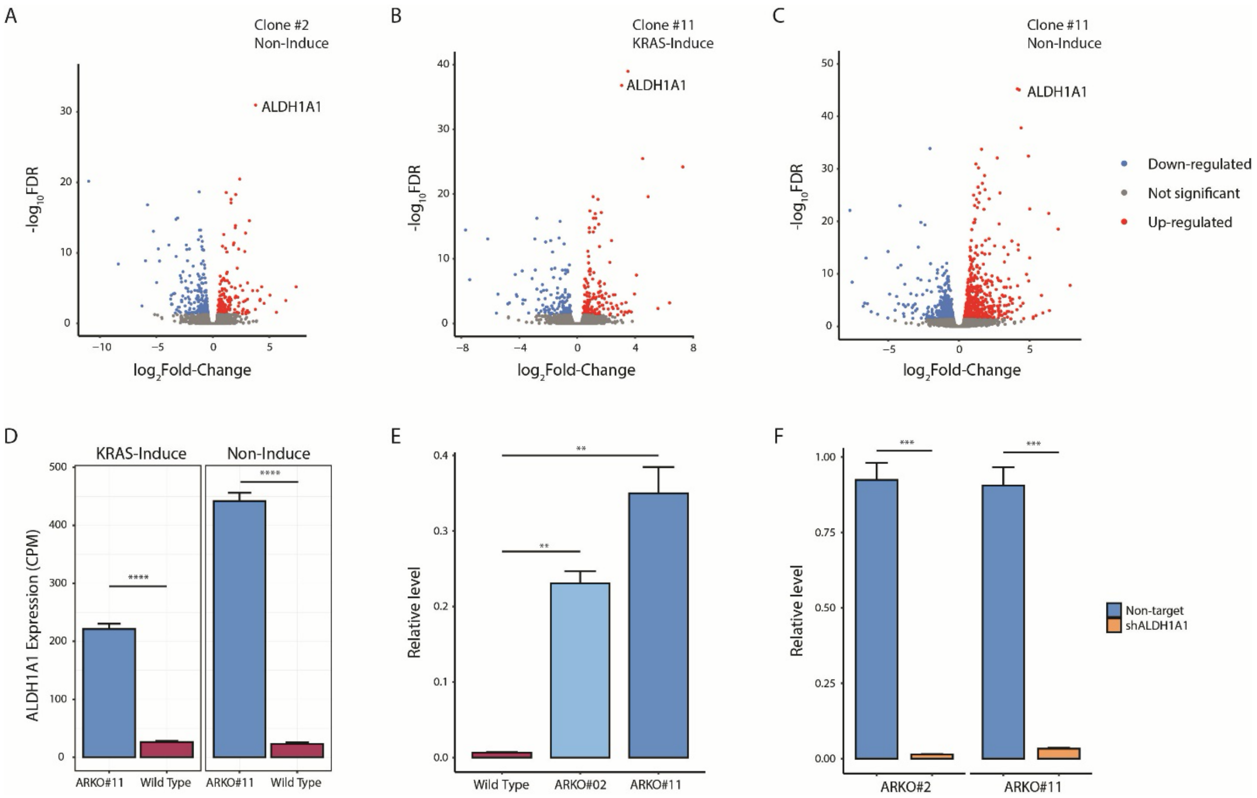


**Figure S6.** *ALDH1A1* expression in *ARID1A*-KO HPNE cells. (**A**) Volcano plot of differentially expressed genes between *ARID1A*-KO cells (clone #2) and wild type cells without KRAS induction. (**B-C**) Volcano plot of differentially expressed genes between *ARID1A*-KO cells (clone #11) and wild type cells with (**B**) or without (**C**) Kras induction. (**D**) ALDH1A1 mRNA level in *ARID1A*-KO cells (clone #11) and wild type cells with (left) or without (right) Kras induction quantified by RNA-seq. CPM, count per million reads. (**E**) The upregulation of ALDH1A1 expression in HPNE cells with ARID1A-KO (clone #2 and #11) was confirmed by qRT-PCR. (**F**) The knockdown efficiency of ALDH1A1 in HPNE cells with ARID1A-KO (clone #2 and #11) was confirmed by qRT-PCR. Student's t-test: **, $p<0.01$; ***, $p<0.001$, ****, $p<0.0001$.

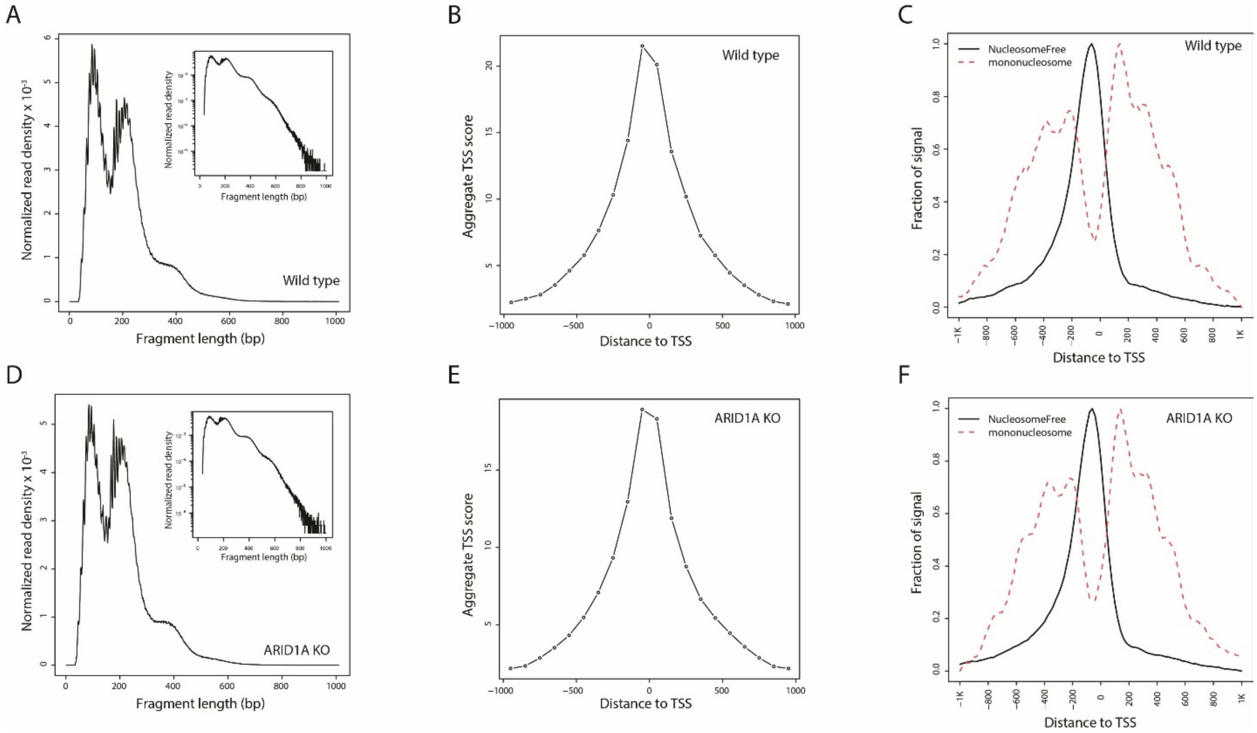


**Figure S7.** Quality control of ATAC-seq. (**A**) The distribution of fragment size of wild type cells. (**B**) The transcription start site (TSS) enrichment score of wild type cells. (**C**) The enrichment of nucleosome-free reads at transcription start sites compared to the reads spanning mono-nucleosome in wild type cells. (**D**) The distribution of fragment size of *ARID1A*-KO cells. (**E**) The TSS enrichment score of *ARID1A*-KO cells. (**F**) The enrichment of nucleosome-free reads at transcription start sites compared to the reads spanning mono-nucleosome in *ARID1A*-KO cells.

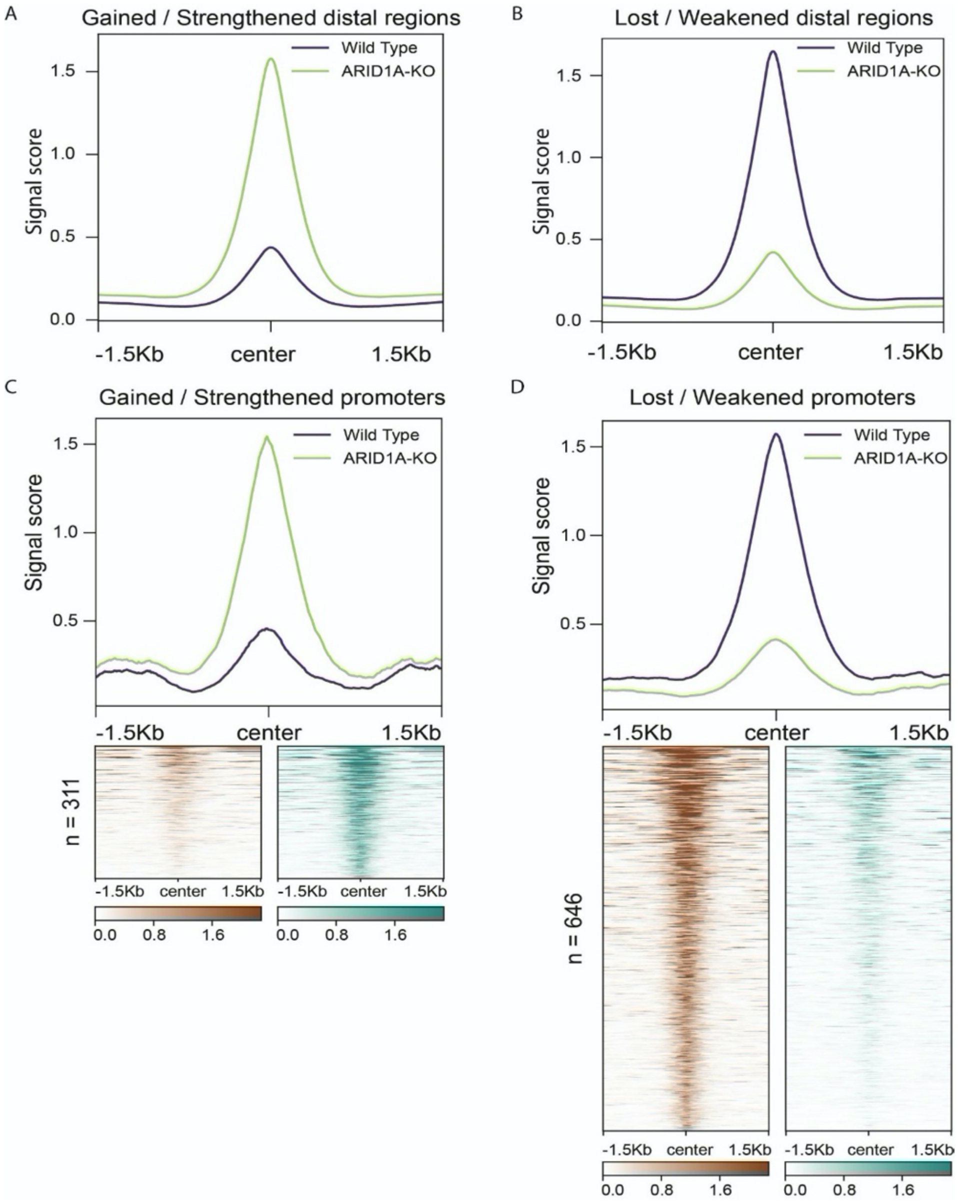


**Figure S8.** Read density profiles of differential peaks. (**A**) The read density profiles of the distal regions with increased accessibility in *ARID1A*-KO cells compared to wild type cells. (**B**) The read density profiles of the distal regions with decreased accessibility in *ARID1A*-KO cells compared to wild type cells. (**C**) The read density profiles of the promoters with increased accessibility in *ARID1A*-KO cells compared to wild type cells. (**D**) The read densities profiles of the promoters with decreased accessibility in *ARID1A*-KO cells compared to wild type cells.

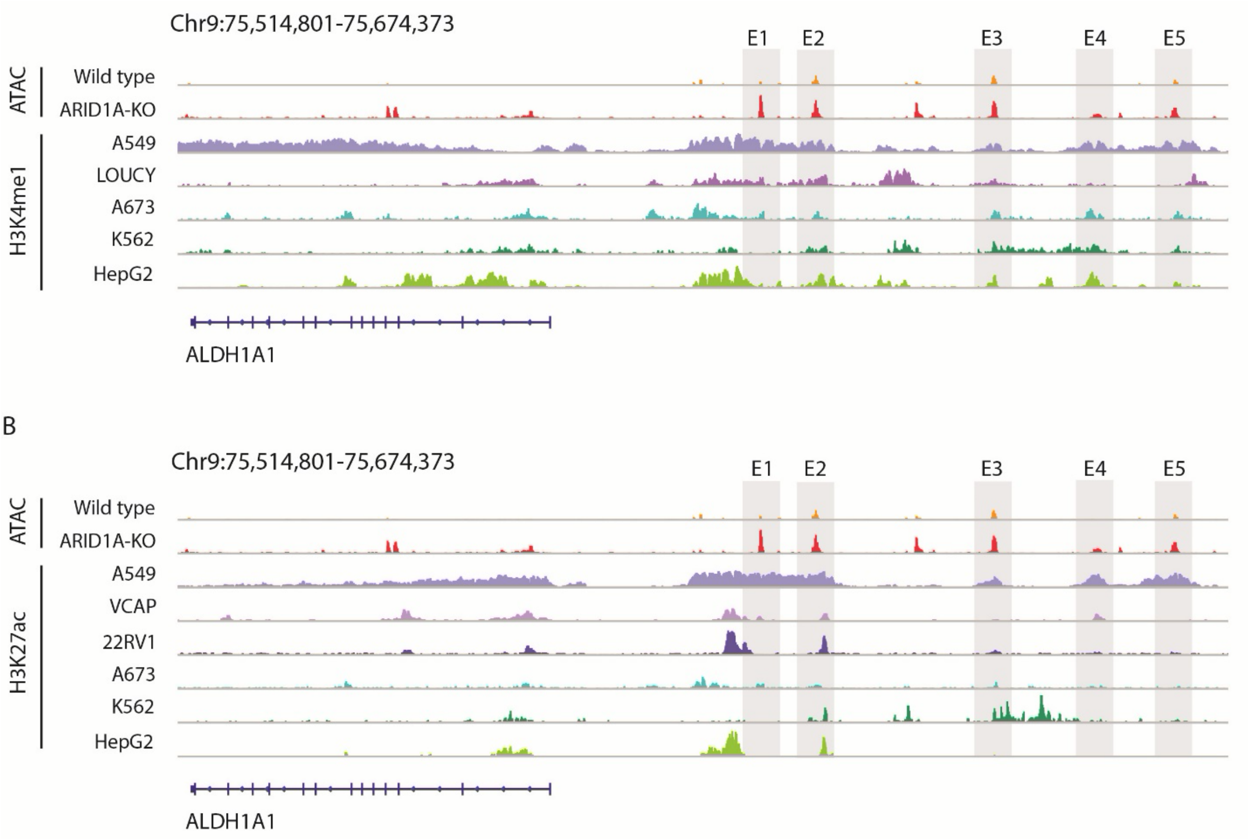


**Figure S9.** The landscape of H3K27ac and H3K4me1 at the upstream of the *ALDH1A1* gene in ALDH1A1$^{high}$ cell lines. (**A**) The landscape of H3K4me1 at the upstream of the *ALDH1A1* gene in five ALDH1A1$^{high}$ cell lines. (**B**) The landscape of H3K27ac at the upstream of the *ALDH1A1* gene in six ALDH1A1$^{high}$ cell lines.

| Peak | Potential binders |
|---|---|
| **Enhancer 1** (chr9:75602590-75603348) | **EP300**, **NR3C1**, **CEBPB** |
| **Enhancer 2** (chr9:75610523-75611906) | NA |
| **Enhancer 3** (chr9:75637758-75638714) | **CREB1**, **SRF**, **ZBTB33** |
| **Enhancer 4** (chr9:75653198-75654412) | **NR3C1**, **NR2F1**, **AR** |
| **Enhancer 5** (chr9:75664913-75666233) | **EP300**, **ZBTB40**, **RELA**, **NR3C1** |

**Figure S10.** The list of TFs that can potentially bind the enhancer regions based on 7 ALDH1A1$^{high}$ cell lines and 5 ALDH1A1$^{low}$ cell lines from the ChIP-seq dataset (Cistrome DB). For the TFs whose binding events are preferentially detected in the datasets from ALDH1A1$^{high}$ cell lines (Fisher's test, $p < 0.05$), we marked them in red color. For the TFs that do not have enough datasets for a statistical test, we marked the TFs whose binding events are observed in more than 50% of ALDH1A1$^{high}$ cell lines in blue color; and the TFs whose binding events are observed in more than 25% but less than 50% of ALDH1A1$^{high}$ cell lines in black color. The TFs that are not expressed in HPNE cell lines are removed.
